# Large-scale experimental removal of non-native slider turtles has unexpected consequences on basking behavior for both conspecifics and a native, threatened turtle

**DOI:** 10.1101/312173

**Authors:** Max R. Lambert, Jennifer M. McKenzie, Robyn M. Screen, Adam G. Clause, Benjamin J. Johnson, Genevieve G. Mount, H. Bradley Shaffer, Gregory B. Pauly

## Abstract

The red-eared slider turtle (*Trachemys scripta elegans*; RES) is one of the world’s most invasive species. Native to the central United States, RES are now widely established in freshwater habitats across the globe, largely due to release of unwanted pets. Laboratory and mesocosm experiments suggest that introduced RES are competitively dominant to native turtles, but such competition remains untested in the wild. Here, we experimentally removed introduced RES to test whether they compete for critical basking habitat with native, threatened western pond turtles (*Emys marmorata*; WPT), a species being considered for listing under the U.S. Endangered Species Act. Following removal, we found that both the remaining RES as well as WPT altered their basking distribution but in a manner inconsistent with strong interspecific competition. However, these findings suggest strong intraspecific competition for basking sites amongst RES and that interspecific competition between WPT and introduced RES likely occurs at higher RES densities. Our works suggests RES influence the behavior of native species in the wild and indicates that RES removal may be most beneficial at high RES densities. This experiment highlights the importance of considering experimental venue when evaluating competition between native and non-native species and should encourage conservation biologists to treat removal efforts as experiments.

## 1.0 Background

Invasive species are a major threat to biodiversity (Simberloff et al. 2013) and are an ongoing concern for conservation practitioners (Kuebbing and Simberloff 2015). One species widely considered harmful to native species worldwide is the red-eared slider turtle (*Trachemys scripta elegans*; RES). This species is native to the central United States but is now present on every continent except Antarctica, predominantly because of releases of unwanted pet turtles (Arvy 1997, Cadi et al. 2008, Kraus 2009, Rhodin et al. 2017). The widespread continued introduction of this species led the International Union for Conservation of Nature (IUCN) to name RES as one of the “worst invasive species” in the world (Lowe et al. 2000). However, despite long-held concerns about the effects of introduced RES on native turtle species (Arvy and Servan 1998, Cadi et al. 2008), few studies have explicitly explored the consequences of RES introductions on wild, native turtle populations (Lambert et al. 2013, Pearson et al. 2013, Costa 2014, Héritier et al. 2017) and there have been no experiments on wild populations.

Laboratory and mesocosm experiments suggest that RES can outcompete native turtles for food and basking sites (Cadi and Joly 2003, 2004, Polo-Cavia et al. 2008, 2010, 2011, Pearson et al. 2015). While these simplified, semi-natural experiments allow us to begin isolating causal agents, they also frequently inflate the effects of interspecific competition compared to *in situ* manipulations under more natural conditions (Skelly and Kiesecker 2001, Skelly 2002, Winkler and Van Buskirk 2012). Although we recognize that *in situ* experiments come with their own drawbacks, comparing laboratory and mesocosms experiments with field manipulations is critical to understanding the strength of species interactions in wild contexts. To our knowledge, no study has yet experimentally tested whether RES are an important competitor with any native turtle species in the wild.

Basking sites are a key resource for evaluating competition between aquatic turtle species because these sites are critical for proper thermoregulation, which directly influences vital physiological parameters like disease control as well as growth and reproductive rates (Ernst and Lovich 2009). Basking sites have repeatedly been identified as a likely axis of competition between introduced RES and native turtle species, with several laboratory and mesocosm experiments suggesting that RES may exhibit dominant aggressive behaviors while basking and may displace native turtles from basking sites (Cadi and Joly 2003, Polo-Cavia et al. 2010, Pearson et al. 2015). In human-modified waterways, competition for basking sites may be especially pronounced because turtles often experience reductions in basking site availability due to the removal of basking objects for flood control and aesthetic reasons (Spinks et al., 2003).

One study in the University of California, Davis (UCD) Arboretum waterway found that RES and native western pond turtles (*Emys marmorata*; WPT) are spatially segregated across basking sites (Lambert et al. 2013). Although both RES and WPT sometimes bask at the same sites (Fig. 1), they tended to concentrate in opposite ends of the waterway and at basking sites that differ in slope, water depth adjacent to the site, site substrate, and the degree of human activity (Lambert et al. 2013). It is unclear, however, whether these interspecific differences in basking site use are due to innate preferences or competitive interactions. Because of the biological importance of basking sites in WPT life history (Bury and Germano 2008, Ernst and Lovich 2009), determining whether RES limit WPT use of preferred basking habitat is essential for effective conservation (Thomson et al. 2016), particularly given the widespread occurrence of introduced RES in California (Thomson et al. 2010, Fisher unpubl.)

**Figure 1.**
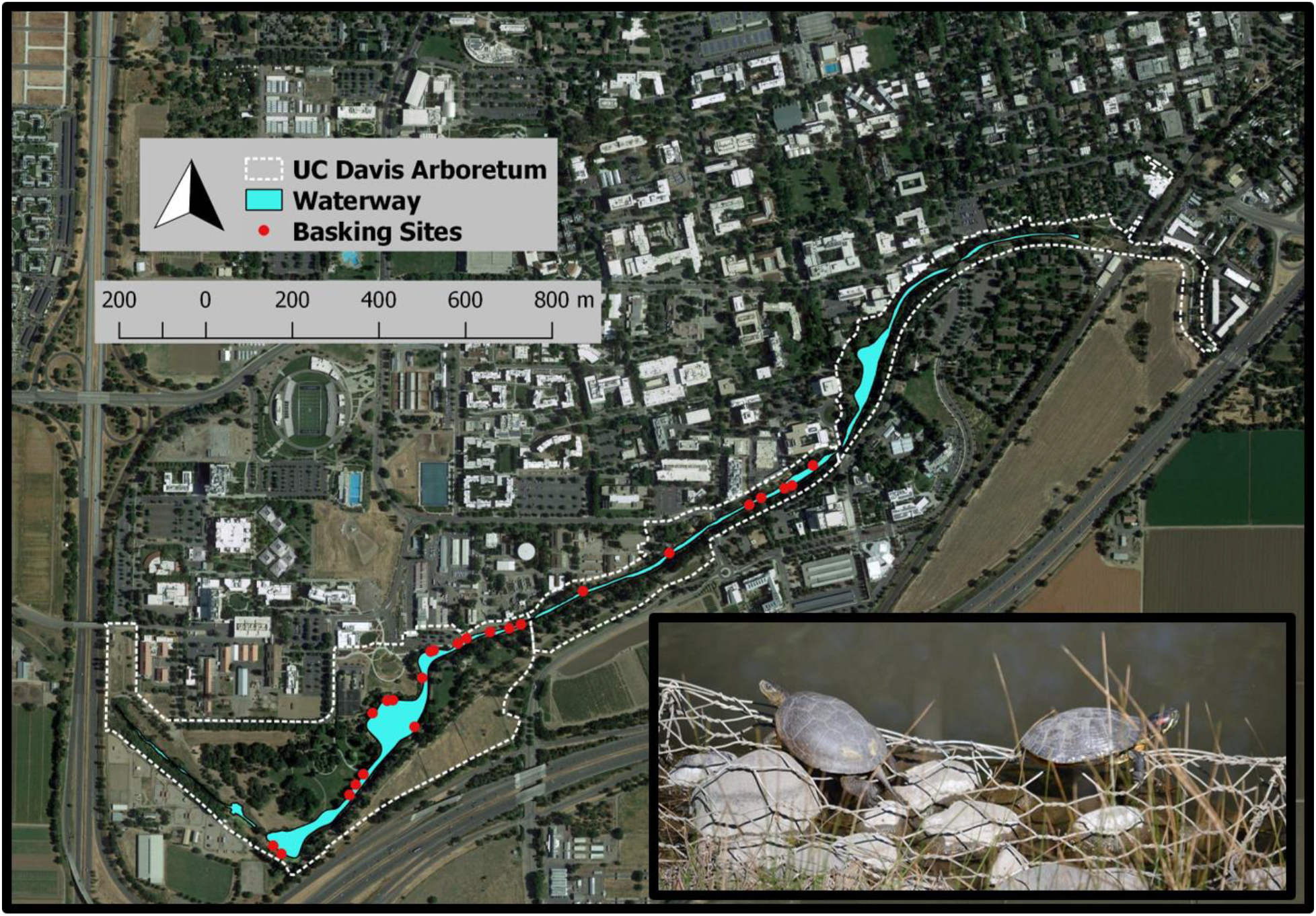
(1.5 columns, color online only): Map of the UC Davis Arboretum waterway and turtle basking sites monitored before and after the red-eared slider population reduction. Inset are a native western pond turtle (left) and a non-native red-eared slider (right) basking in the UC Davis Arboretum.Photo by M. Lambert.

Here, we present the results of an *in situ* field experiment whereby we dramatically reduced the UCD Arboretum RES population to examine whether WPT subsequently shifted their use of available basking sites in the wild. Our experiment explicitly tests whether invasive species removal, an intensive and commonly-advocated management practice (Simberloff et al. 2013, Gaeta et al. 2015), including for RES (Garcia-Díaz et al. 2017), influences the basking behavior of native WPT in the wild. If RES and WPT compete for basking sites and RES are dominant to WPT, then we predict that removing RES would lead to WPT basking activity becoming more concentrated at sites previously dominated by RES. However, if existing basking-site use patterns reflect species-specific habitat preferences, then we predict that removing RES would have minimal impact on WPT basking site use. Results from this experiment provide a useful first test of the impacts of introduced RES on native, wild turtles; these data are immediately relevant to management of WPT across its known range (Thomson et al. 2016), and for the undergoing Status Review for possible listing under the US Endangered Species Act (USFWS 2015)

## 2.0 Methods

### 2.1 Study Site

The UCD Arboretum waterway runs along the southern border of the university campus in Yolo County, CA, USA and is situated in the former channel of the North Fork of Putah Creek. Various sections of the waterway are bordered by urban, agricultural, and undeveloped natural landscapes (Fig. 1). For more detailed descriptions of the location, see Spinks et al. (2003) and Lambert et al. (2013).

### 2.2 RES Removal

In 2011, we captured turtles throughout the UCD Arboretum from 10 July–1 August and again from 13–29 September. We primarily used baited traps that can be deployed in water depths of 0.5–2.0 m. Cumulative submersible trap effort was approximately 900 trap-nights. We supplemented our submersible trapping with opportunistic hand captures and dip netting, along with periodic deployment of a fyke net and a basking trap. The submersible traps, fyke net, and basking trap were not biased towards any particular species, but hand captures and dip netting were targeted at RES. We recorded mass and plastron length of each captured RES using digital pan scales and dial calipers. We re-homed several captured RES with responsible pet owners and euthanized all other RES, donating the majority to the UC Davis School of Veterinary Medicine, the Natural History Museum of Los Angeles County, or the Museum of Wildlife and Fish Biology at UC Davis. All turtle handling was authorized under UC Davis IACUC Protocols #15263 and #16227, and California Department of Fish and Wildlife Scientific Collecting Permits #2480, #4307, and #11663.

To test whether our RES trapping success plateaued over time, which would suggest that our trapping effort removed the majority of the RES population, we analyzed whether the cumulative number of RES was better modeled by a linear or quadratic relationship across trapping days. We used linear regression and likelihood ratio tests to determine whether our trapping effort had minimal impact on the RES population (a linear fit) or resulted in fewer RES trapped each day (a quadratic fit). We also tested for an interaction between sex and trapping day to estimate whether we reduced the sexes at different rates.

### 2.3 Turtle Monitoring

From 18 March–22 April 2012, we conducted visual (with binoculars) surveys of the same set of 24 basking sites studied in spring 2010 prior to the RES removal (Fig. 1). Each basking site is a short stretch of shoreline (1–2 m long) with adjacent sites at least 3 m apart. We conducted surveys 2010 and 2012 surveys within a similar set of dates and times to make them as similar as possible. During each survey, we measured water temperature with a hand-held thermometer; we also obtained maximum daily temperature data from the UC Davis Russel Ranch Weather Station, ca. 4 km northwest of the UCD Arboretum.

### 2.4 Analysis

The basking distribution of WPT and RES previously was shown to vary strongly along a west-east gradient, with WPT focused at the west end and RES focused at the east end of the waterway (Lambert et al. 2013). Because of this, we analyzed the relative and absolute basking abundances of both species, and the extent to which these changed after removing RES. We limited our analysis to 24 basking sites that had data available for every survey date within the same date range (March 18 to April 22) in 2010 (pre-RES removal, from Lambert et al. 2013) and in 2012 (post-RES removal, measured here).

To test for changes in the relative basking distribution of WPT to RES across the waterway, we used a generalized linear mixed effects model (GLMM) with a binomial family for proportion data using the ‘glmer’ function in the R package lme4. We used the distance of each basking site from the west end of the UCD Arboretum (following Lambert et al. 2013) as well as treatment (pre- or post-RES removal) as fixed effects and used observation date as a random effect to account for repeated measures of basking sites (Lambert et al. 2013). We first tested for a significant interaction between treatment and each basking site’s distance from the west end. If the interaction term was not significant, we removed it from the model. We then assessed the influence of the main effects and tested whether the relative basking distribution of WPT to RES differed pre- and post-RES removal using a Tukey’s post-hoc test with ‘glht’ function in the R package “multcomp”. We also performed binomial GLMMs for each basking site separately to explore whether individual basking sites show changes in the proportion of WPT to RES after the experiment.

For the absolute abundance of each species, we applied a similar modeling approach but used Poisson GLMMs for count data. We used the ‘r.squaredGLMM’ function in the package “MuMIn” to calculate R^2^ for GLMMs; ‘MuMIn’ calculates both a conditional R^2^ (cR^2^) for the full model including fixed and random effects as well as a marginal R^2^ (mR^2^) for just the model’s main effects. We conducted all analyses in R (version 3.2.2).

To further explore patterns at individual basking sites, we used contingency table analyses for each species independently to test whether certain basking sites comprised larger or smaller proportions of the total basking observations for either species pre- and post-RES removal. We focused on sites P, O, E, Q, and R which were, respectively, the five most heavily-used turtle basking sites (combined for both species) pre-RES removal. We also examined site X since it was the most heavily-used turtle basking site post-RES removal.

## 3.0 Results

### 3.1 RES Removal

In summer 2011, we captured and removed 180 RES from the UCD Arboretum. We removed 25 adult males, 71 adult females, and 84 juveniles (individuals with carapace length ≤ 100 mm, Ernst and Lovich 2009), including one individual of 113 mm carapace length that lacked sexually-diagnostic traits. We removed 59 RES that we had captured and marked in previous years and 121 unmarked individuals. All turtle marking previously occurred as part of the UC Davis Herpetology course from 2007 through spring 2011. Of these new RES, 70% (n = 84) were juveniles, 22% (n = 27) were adult females, and 8%(n = 10) were adult males.

A likelihood ratio test supported a model (full model R^2^ = 0.95, p < 0.0001) with a quadratic over a linear fit between cumulative RES trapped and trapping day (p < 0.0001) and with an interaction between RES sex and trapping day (p < 0.001). During the removal effort, captures of RES declined and leveled off, indicating that we removed a substantial portion of the catchable RES population. Furthermore, the significant interaction term suggests we depleted the male RES population faster than the female RES population (Fig. 2). In total, we removed 104.5 kg of RES biomass, of which 79% (82.3 kg) was from adult females, 15% (15.5 kg) was from adult males, and 6% (6.7 kg) was from juveniles. During this same trapping effort, we captured, marked (or re-marked), and released 118 individual WPT, 14 of which were juveniles (≤ 110mm plastron length; Holland, 1991; Spinks et al. 2003). While some aspects of our capture efforts in 2011 specifically targeted RES (e.g., dip netting and hand captures), our data indicate RES outnumbered WPT by about 1.5:1 at the start of the experiment.

**Figure 2.**
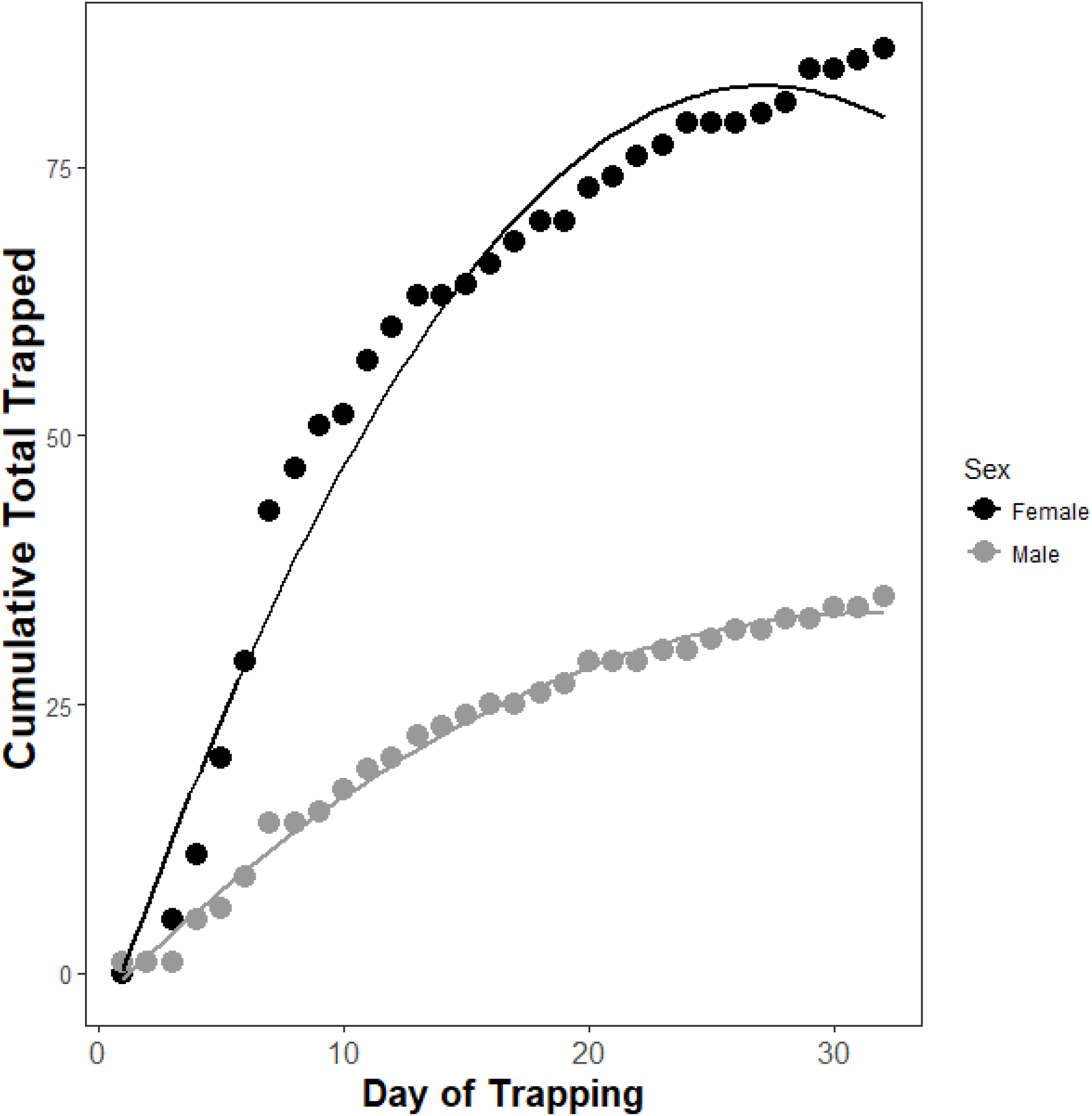
(single column). Cumulative total number of adult female and male red-eared sliders (RES; *Trachemysscriptaelegans*) removed from the UC Davis Arboretum in 2011. Trap Day 1–18 are in July and 19–32 are in September.

### 3.2 Basking Surveys

We surveyed for 16 days from 18 March to 22 April 2010 (pre-removal) and 18 days from 18 March to 22 April 2012 (post-removal). Maximum daily air temperatures were not significantly different between years (two-tailed t-test, p = 0.74; 2010, 19.2 C ± 0.69 SE; 2012, 18.8 C ± 0.88 SE). However, in the two weeks prior to our surveys the maximum daily air temperatures were significantly warmer in 2012 (18.8 C ± 1.08 SE) than in 2010 (15.2 C ± 0.65 SE); two-tailed t-test, p < 0.001). Water temperature was significantly warmer (two-tailed t-test, p < 0.0001) in 2010 (17.0 C ± 0.24 SE) compared to 2012 (15.4 C ± 0.36 SE). In 2010, we recorded 283 WPT and 645 RES observations. In 2012, we recorded only 43 WPT observations and 61 RES observations.

Pre-removal, we recorded WPT basking at 15 of the 24 basking sites, but post-removal we recorded WPT basking at only 8 of the 24 sites (Fig. 3). WPT were absent from eight sites that they used pre-removal, although six of these were used infrequently in 2010. We recorded WPT using one additional site where they were not recorded pre-removal. In general, the basking sites most commonly used by WPT pre-removal were the same sites used post-removal (Fig. 3). Pre-removal, we recorded RES basking at 17 of the 24 basking sites, but post-removal we recorded RES basking at only 8 of the 24 sites (Fig. 3). RES were absent from nine sites that they used pre-removal and were not recorded using any new sites after the removal.

**Figure 3.**
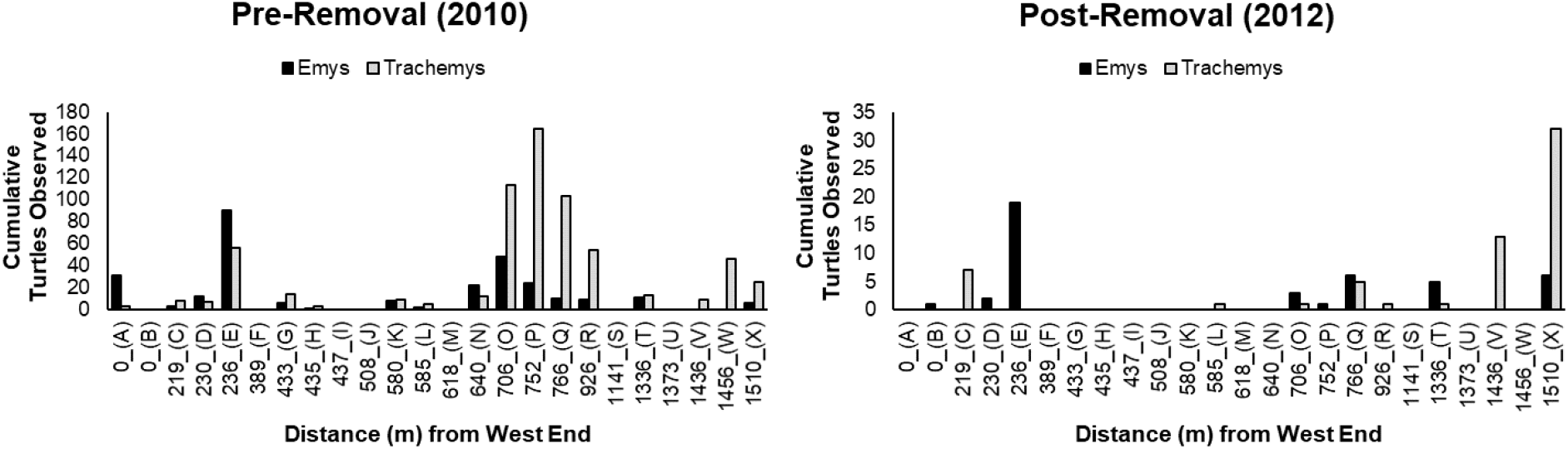
(double column). The cumulative number of western pond turtle (WPT; *Emys marmorata*) and red-eared slider (RES; *Trachemysscriptaelegans*) basking observations across sampling dates in the pre- and post-RES removal years. Letters in parentheses under the x-axis are basking site identifiers. Note that y-axes of the two panels have different scales.

### 3.3 Relative Abundance

The interaction between distance from the west end of the waterway and treatment was not significant (p = 0.18) and was removed from the model. Both distance from the west end (p < 0.0001) and treatment (p < 0.0001) were significant and were retained in the model (cR^2^ = 0.31, mR^2^ = 0.31). Both pre- and post-RES removal, the relative basking distribution of turtles was WPT-biased in the west end and RES-biased in the east end of the waterway (Fig. 4). Furthermore, the Tukey’s post-hoc test indicated that the proportion of basking observations increased from 30.5% WPT pre-RES removal to 41.3% WPT post-RES removal (p < 0.0001). The non-significant interaction term indicates that the RES removal did not change the relative basking distribution of the two species across the waterway.

**Figure 4.**
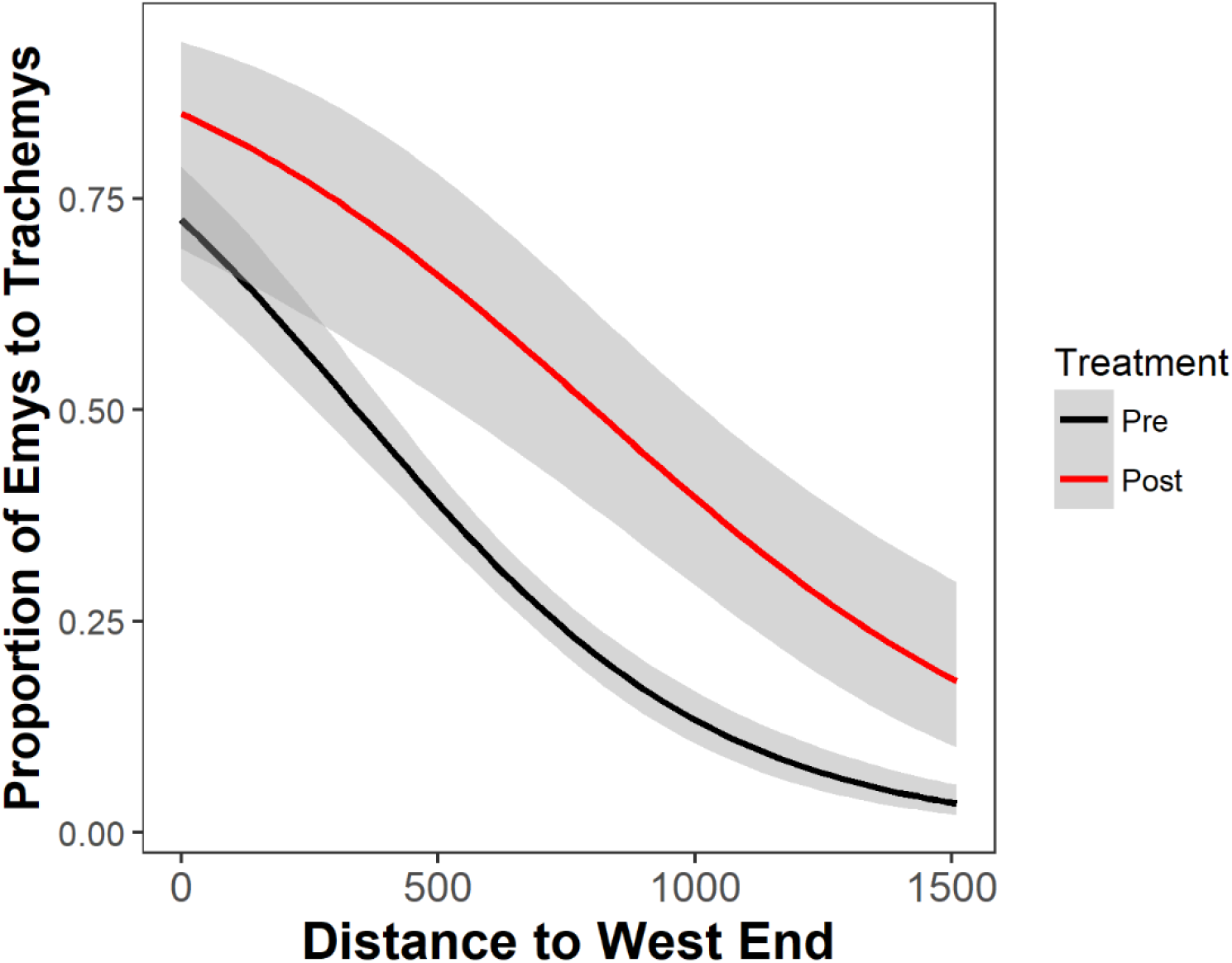
(single column, color in print). The modeled proportion of western pond turtles (WPT; *Emys marmorata*) to red-eared sliders (RES; *Trachemysscriptaelegans*) basking along a west-east gradient in the UC Davis Arboretum. While WPT made up a greater proportion of observations in 2012 than in 2010, the relative basking distribution of the two species along the Arboretum did not change between pre- and post-RES reduction years. Curves are binomial fits and gray shading represents 95% confidence intervals.

Individual binomial GLMMs for each basking site returned a significant treatment effect for site Q (p = 0.002, 9% WPT to 55% WPT) and a marginally significant effect for site O (p = 0.09, 30% WPT to 75% WPT). All other basking sites showed no significant difference in the proportion of the two species between years (all p > 0.1).

### 3.4 WPT Absolute Abundance

In 2010, we recorded 283 WPT basking observations and only 43 in 2012. The Poisson GLMM indicated a significant interaction between treatment and distance from the west end (p = 0.012, cR^2^ = 0.23, mR^2^ = 0.06), suggesting a shift in the absolute basking distribution of WPT across the waterway. Individual GLMMs for each year indicate that distance from the west end is significant in the pre-RES removal year (p < 0.012, cR^2^ = 0.27, mR^2^ = 0.03) but not in the post-RES removal year (p = 0.55). These results suggest that, before the RES removal, absolute basking abundance of WPT declined from west-east, and that post-RES removal WPT had a relatively even basking distribution throughout the UCD Arboretum (Fig. 5). Contingency tables indicated that sites Q (p = 0.01) and X (p = 0.001) comprised larger proportions of total WPT basking observations post-RES removal than pre-RES removal. All other sites analyzed comprised similar proportions of total WPT basking observations before and after the experiment (all p > 0.1), although small sample sizes often resulted in relatively little statistical power. Together, these analyses indicate removing RES resulted in a less clustered, more even distribution of WPT across basking sites with two sites towards the center-east and east of the Arboretum comprising more WPT basking activity.

**Figure 5.**
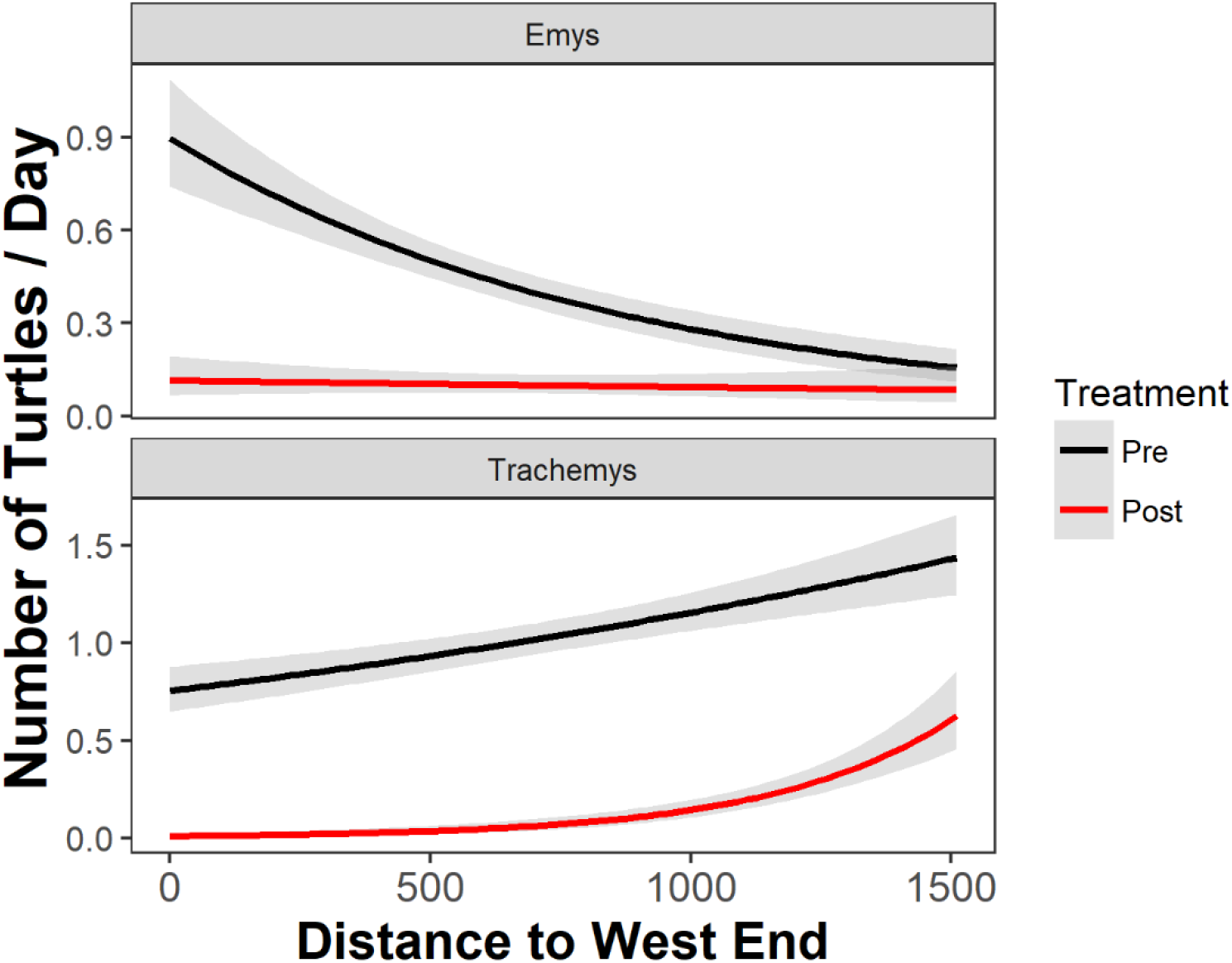
(single column, color in print). The modeled number of western pond turtles (WPT; *Emys marmorata*) and red-eared sliders (RES; *Trachemysscriptaelegans*) observed basking along a west-east gradient at the UC Davis Arboretum pre- and post-RES removal. Note that fewer turtles were observed basking in the post-RES removal survey. Curves are Poisson fits and gray shading represents 95% confidence intervals.

### 3.5 RES Absolute Abundance

For RES, the GLMM indicated a significant interaction between distance to the west end and treatment (p < 0.0001, cR^2^ = 0.30, mR^2^ = 0.16). While the number of RES basking observations was an order of magnitude lower in 2012 (n = 61) versus 2010 (n = 645), the positive relationship between RES absolute abundance and the west-east gradient in the UCD Arboretum appears to be more pronounced post-RES removal (Figs. 3, 5). Individual GLMMs for each year show that distance to the west end was significant in the pre-RES removal year (p < 0.0001, cR^2^ = 0.27, mR^2^ = 0.02) and the post-RES removal year (p < 0.0001, cR^2^ = 0.14, mR^2^ = 0.14). In both years, the absolute abundance of basking RES increased along the west-east gradient pre (Fig. 5). After the RES-removal, RES were relatively sparse through the western and central portions of the waterway and were concentrated in the far eastern end (Fig. 3, 5). Contingency table analyses indicated that sites E (p = 0.01), O (p = 0.0004), P (p = 0.0001), and R (p = 0.03) comprised lower proportions of total basking observations after the experiment and site X comprised a higher proportion (p = 0.001). Site Q made up similar proportions of total RES observations in both years (p = 0.14).

## 4.0 Discussion

To test whether non-native RES influence WPT basking site use, we removed 180 non-native RES, totaling over 100 kg of turtle biomass. This experiment represents a dramatic alteration to the turtle community inhabiting the UCD Arboretum. Our experiment indicated that removing the majority of the RES population altered the basking distribution of both native WPT and residual RES, and that some form of interspecific interactions is occurring between the two species. However, our results do not necessarily provide evidence for strong interspecific competition between introduced RES and WPT, and suggest that a more nuanced, complex set of interactions may be occurring in wild populations.

### 4.1 Intraspecific Competition

One of the clearest effects of our experiment was that the remaining post-removal RES abandoned several basking sites that they previously used heavily (particularly sites O and P) and shifted towards the east end of the transect (e.g., sites V and X). Although it is unclear what drove this shift, this result indicates that RES prefer habitat at this end of the waterway and that, prior to our experiment, RES densities were high enough for intraspecific competition among RES to force some individuals into other areas of the waterway. Our previous work showed that RES basking activity was highest at sites with shallow slopes, deeper water adjacent to the site, a steel mesh (rather than concrete or dirt) substrate, and high human activity (Lambert et al. 2013). Consistent with these observations, the two basking sites that showed the most concentrated RES activity post-removal were comprised of steel mesh and were the sites with some of the flattest slopes and deepest water along the transect as well as the sites with the highest level of human activity (Lambert et al. 2013), indicating that residual RES concentrated their basking activity at the most preferred sites.

### 4.2 Interspecific competition

Previous work on the UCD Arboretum turtle population found that RES and WPT largely use different basking sites (Lambert et al. 2013). Before and after our experiment, WPT predominantly used the same basking sites but at different frequencies, with a general trend toward a more uniform west-east distribution post-removal. These results indicate that reducing the density of introduced RES allow WPT to spread out in the waterway. Even so, if sites towards the east end of the waterway which were previously dominated by RES (e.g., sites O, P, Q, and R) are also preferred by WPT, then we would have expected WPT basking behavior to concentrate at these sites post-RES removal. But we did not see this. Rather, our experiment resulted in a shift in WPT basking activity suggesting that WPT basking is contingent on RES densities but we did not observe a dramatic shift that might be indicative of strong interspecific competition for basking sites. Competition is presumably greatest at high densities of RES and perhaps influenced by the relative densities of both species, as has been shown in other biological invasion scenarios (Gurnell et al. 2004). Earlier experiments have concluded that introduced RES outcompete native turtles for resources including basking sites or food. Because these experiments took place in artificial experimental venues and (Cadi and Joly 2003, 2004, Polo-Cavia et al. 2008, 2010, 2011, Pearson et al. 2015), it is possible that prior conclusions about the competitive dominance of introduced RES were inflated. Although no prior experiments have focused on WPT, some (Cadi and Joli 2003, 2004) have focused on European *Emys orbicularis*, which is a closely-related congener (Spinks et al. 2016).

### 4.3 Study Limitations

Our analyses showed that WPT made up proportionally more of our observations after RES removal (Fig. 4). However, we recorded far fewer turtle basking observations for both species in 2012. For RES, this was expected as it was the goal of our experiment to reduce the RES population. This is not the case for WPT. It is possible that temperature or other environmental variation among years as well as unforeseen consequences of our manipulation resulted in reduced overall turtle basking activity after the RES removal. For instance, aquatic turtles like RES can dramatically influence trophic dynamics and aquatic ecosystem function (Lindsay et al. 2013). By removing a substantial portion of the turtle community, our experiment may have altered the availability and distribution of food resources which may have indirectly impacted where turtles chose to bask and turtle basking behavior generally, resulting in fewer observations. Unfortunately, we cannot distinguish whether the generally lower basking observations of WPT post-RES removal are an effect of our experiment or whether other uncontrolled factors may have resulted in fewer WPT basking observations. Because of logistical constraints, we were only able to collect a single of year of observations post-RES removal. We also recognize that our experiment did not address other putative axes of competition that are important for the continued recruitment and persistence of this WPT population. For example, evidence from experimental mesocosms suggests introduced RES generally eat more and grow faster than native turtles (Cadi and Joly 2003, Pearson et al. 2015), and these effects may have important consequences for native turtles.

### 4.4 Management Implications

Removing 180 RES from the UCD Arboretum was an intensive effort requiring over 2,000 person-hours of field work across forty days. Although WPT comprised a larger proportion of our basking observations post-removal (Fig. 4), RES still made up the dominant portion of our observations, summing to almost 60% of the total basking observations made after the experiment. In general, removing invasive species is difficult, time and labor intensive, and may still fail to extirpate the entire population, particularly in the face of continued introductions (Gaeta et al. 2015). In Europe, where RES removal is a widely advocated practice, recent work noted the severe challenges of functionally eradicating introduced RES (Garcia-Diaz et al. 2017). As long as RES are readily available in the pet trade, *de novo* introductions are likely to continue, complicating attempts to successfully eradicate introduced RES populations.

Our results suggest that a concerted effort at RES reduction in a large, complex water body has the potential to influence native turtle species, but that these influences may be relatively modest in their quantitative effects. Regardless of whether it is known that RES compete with a given native species, both removing non-native RES and stemming the future release of RES are important steps for reducing possible disease and parasite transmission (Héritier et al. 2017; Demkowska-Kutrzepa et al. 2018). Further, removals and reductions in pet releases could help minimize competition if it is occurring, whether that be for food, basking sites, or other resources. Although the commitment of time and energy is large, we encourage conservation biologists to treat RES removal efforts as experiments, as was done here, and test whether removing RES benefits native turtle species along these other ecological axes.

Habitat modification due to urban and agricultural land use is a major threat to WPT in California (Thomson et al. 2016). Nonetheless, human-modified habitats can be valuable resources for WPT when appropriate conservation and management efforts are implemented (Spinks et al. 2003, Thomson et al. 2010, 2016). Directly managing urban basking habitat may be a particularly tractable conservation activity for WPT in addition to directly managing non-native RES. Future experiments and management practices can readily manipulate these basking site characteristics to test whether doing so is beneficial for WPT. Emerging research from the UCD Arboretum suggests that experimentally-added floating logs are preferred by WPT over bank-side basking and are more heavily used by WPT than RES especially when placed further from human activities (Cossman et al. unpubl.). In our experiment here, we may have liberated parts of the waterway that were previously dense with turtles generally, thus allowing WPT to spread out across the waterway. However, because WPT did not concentrate their basking activity at sites previously dominated by RES, these two species may not intensely compete for bank-side basking sites in this waterway. WPT may ultimately show little preference for particular bank-side basking site characteristics, although this warrants further study. Providing more basking sites of suitable quality, and particularly further from high levels of human activity, may be a feasible and fruitful management practice in conjunction with removing RES. We encourage additional research into the merits of this strategy.

### 4.5 Conclusions

Evidence from laboratory and mesocosm studies indicates that introduced RES are competitively dominant to native turtles. Here, we offer the first experimental test for competition between native turtles and non-native RES in the wild; our work provides insight into the seemingly complex nature of competition between introduced RES and native turtles. Our population manipulation suggests that reducing the density of RES may alter the basking activity of threatened WPT but that RES and WPT may not compete intensely for basking sites. We found strong evidence for strong intraspecific competition for basking sites at high RES densities, and that reducing that competition may have had additional effects on the distribution of WPT basking. We hope that our study will encourage further field-based experiments to better understand the extent to which RES are competing with native turtles for basking sites and/or other resources, and to explore which management practices are both reasonable and effective.

## Acknowledgments

This work was initiated while all authors were at UC Davis. This research did not receive any specific grant from funding agencies in the public, commercial, or not-for-profit sectors. There are no financial benefits that can arise from this work. We thank Ellen Zagory, Shannon Still, and the staff of the UCD Arboretum and Public Garden for assistance with this experiment and for access to GIS files. Members of the Peter Moyle Lab and Janet Foley Lab provided equipment and supplies for removing RES. Nick Buckmaster, Lauren Cassidy, Jillian Howard, Anna Jordan, Brian Mahardja, Hilary Rollins, Bryce Sullivan, and Matthew Young provided invaluable field assistance. The UCD Arboretum is heavily used by UC Davis affiliates and members of the public; we thank the many Arboretum visitors who were curious about our research and/or who encouraged our activities.

## Literature Cited

Arvy, C. 1997. Le commerce de *Trachemys scripta elegans*: une menace d’expansion de l’espece dans le monde entire. Bulletin de la Societe Herpetologique de France 84:15–24.

Arvy, C., and J. Servan. 1998. Imminent competition between *Trachemys scripta* and *Emys orbicularis* in France. Mertensiella 10:33–40.

Bury, R. B. and D. J. Germano. 2008. *Actinemys marmorata* (Baird and Girard 1852) – western pond turtles, Pacific pond turtles. Chelonian Research Monographs 5: 001.1–001.9.

Cadi, A. and P. Joly. 2003. Competition for basking places between the endangered European pond turtle (*Emys orbicularis galloitalica*) and the introduced red-eared slider (*Trachemys scripta elegans*). Canadian Journal of Zoology 81: 1392–1398.

Cadi, A. and P. Joly. 2004. Impact of the introduction of the red-eared slider (*Trachemys scripta elegans*) on survival rates of the European pond turtle (*Emys orbicularis*). Biodiversity and Conservation 13: 2511–2518.

Cadi, A., Teillac., Delmas, V., Girondot, M., Servais, V., and A-C. Prevot-Julliard. 2008. Slider turtles *Trachemys scripta elegans* released in France: a case of integrated research and conservation program. Revista Española de Herpetología 22: 111–114.

Costa, Z. J. 2014. Responses to predators differ between native and invasive freshwater turtles: environmental context and its implications for conservation. Ethology 120: 1–8.

Demkowska-Kutrzepa, M., M. Studzinska, M. Roczen-Karcmarz, K. Tomczuk, Z. Abbas, and P. Rózanski. 2018. A review of the helminths co-introduced with Trachemys scripta elegans—A threat to European native turtle health. Amphibia-Reptilia IN PRESS.

Ernst, C. H. and J. E. Lovich. 2009. Turtles of the United States and Canada. Johns Hopkins University Press, 2nd Ed. 840 pages.

Gaeta, J. W., Hrabik, T. R., Sass, G. G., Roth, B. M., Gilber, S. J., and M. J. Vander Zanden. 2015. A whole-lake experiment to control invasive rainbow smelt (Actinopterygii, Osmeridae) via overharvest and a food web manipulation. Hydrobiologia 746: 433–444.

Garcia-Diaz, P., Ramsey, D. S. L., Woolnough, A. P., Franch, M., Llorente, G. A., Montori, A., Buenetxea, X., Larrinaga, A. R., Lasceve, M., Alvarez, A., Traverso, J. M., Valdeon, A., Crespo, A., Rada, V., Ayllon, E., Sancho, V., Lacomba, J. I., Bataller, J. V., and M. Lizana. 2017. Challenges in confirming eradication success of invasive red-eared sliders. Biological Invasions 19: 2739–2750.

Gurnell, J., Wauters, L. A., Lurz, P. W. W., and G. Tosi. 2004. Alien species and interspecific competition: effects of introduced eastern grey squirrels on red squirrel population dynamics. Journal of Animal Ecology 73: 26–35.

Héritier, L., A. Valdeón, A. Sadaoui, T. Gendre, S. Ficheux, S. Bouamer, N. Kechemir-Issad, L. Du Preez, C. Palacios, and O. Verneau. 2017. Introduction and invasion of the red-eared slider and its parasites in freshwater ecosystems of southern Europe: Risk assessment for the European pond turtle in wild environments. Biodiversity and Conservation 26:1817–1843.

Holland, D. C. 1991. A Synopsis of the Ecology and Status of the Western Pond Turtle Clemmys marmorata in 1991. Report to National Ecological Research Center. United States Fish and Wildlife Service, San Simeon, CA.

Knapp, R. A., Boiana, D. M., and V. T. Vredenburg. 2007. Removal of nonnative fish results in population expansion of a declining amphibian (mountain yellow-legged frog, *Rana muscosa*). Biological Conservation 135: 11–20.

Krauss, F. 2009. Alien reptiles and amphibians, a scientific compendium and analysis. Springer, Netherland. 563 pp.

Kuebbing, S. E. and D. Simberloff. 2015. Missing the bandwagon: nonnative species impacts still concern managers. NeoBiota 25: 73–86.

Lambert, M. R., Nielsen, S. N., Wright, A. N., Thomson, R. C., and H. B. Shaffer. 2013. Habitat features determine the basking distribution of introduced red-eared sliders and native western pond turtles. Chelonian Conservation and Biology 12: 192–199.

Lindsay, M. K., Zhang, Y., Forstner, M. R. J., and D. Hahn. 2013. Effects of the freshwater turtle *Trachemys scripta elegans* on ecosystem functioning: an approach in experimental ponds. Amphibian-Reptilia 34: 75–84.

Lowe, S., Browne, M., Boudjelas, S., and M. De Poorter. 2000. 100 of the World’s Worst Invasive Alien Species. A selection from the global invasive species database. The Invasive Species Specialist Group (ISSG) of the Species Survival Commission (SSC) of the World Conservation Union (IUCN), 12 pp.

Pearson, S. H., Avery, H. W., Kilham, S. S., Velinsky, D. J., and J. R. Spotila. 2013. Stable isotopes of C and N reveal habitat dependent dietary overlap between native and introduced turtles *Pseudemys rubiventris* and *Trachemys scripta*. PLoS ONE: https://doi.org/10.1371/journal.pone.0062891

Pearson, S. H., Avery, H. W., and J. R. Spotila. 2015. Juvenile invasive red-eared slider turtles negatively impact the growth of native turtles: implications for global freshwater turtle populations. Biological Conservation 186: 115–121.

Polo-Cavia, N., Lopez, P., and J. Martin. 2008. Interspecific differences in responses to predation risk may confer competitive advantages to invasive freshwater turtle species. Ethology 114: 115–123.

Polo-Cavia, N., Lopez, P., and J. Martin. 2010. Competitive interactions during basking between native and invasive freshwater turtle species. Biological Invasions 12: 2141–2152.

Polo-Cavia, N., Lopez, P., and J. Martin. 2011. Aggressive interactions during feeding between native and invasive freshwater turtles. Biological Invasions 13: 1387–1396.

Rhodin, A. G. J., Iverson, J. B., Bour, R., Fritz, U., Georges, A., Shaffer, H. B., and P. P. van Dijk. 2017. Turtles of the World. Annotated checklist and atlas of taxonomy, synonymy, distribution, and conservation status (8th Ed.). Chelonian Research Monographs 7: 1–292.

Skelly, D. K. and J. M. Kiesecker. 2001. Venue and outcome in ecological experiments: manipulations of larval anurans. Oikos 94: 198–208.

Skelly, D. K. 2002. Experimental venue and estimation of interaction strength. Ecology 83: 2097–2101.

Simberloff, D., Martin, J-L., Genovesi, P., Maris, V., Wardle, D. A., Aronson, J., Courchamp, F., Galil, B., Garcia-Berthou, E., Pascal, M., Pysek, P., Sousa, R., Tabacchi, E., and M. Vila. 2013. Impacts of biological invasions: what’s what and the way forward. Trends in Ecology and Evolution 28: 58–66.

Spinks, P. Q., Pauly, G. B., Crayon, J. J., and H. B. Shaffer. 2003. Survival of the western pond turtle (*Emys marmorata*) in an urban California environment. Biological Conservation 113: 257–267.

Spinks, P. Q., Thomson, R. C., McCartney-Melstad, E., and H. B. Shaffer. 2016. Phylogeny and temporal diversification of the New World pond turtles (Emydidae). Molecular Phylogenetics and Evolution 103 85–97.

Thomson, R. C., Spinks, P. Q., and H. B. Shaffer. 2010. Distribution and abundance of invasive red-eared sliders (*Trachemys scripta elegans*) in California’s Sacramento River Basin and possible impacts on native western pond turtles (*Emys marmorata*). Chelonian Conservation and Biology 9: 297–302.

Thomson, R. C., Wright, A. N., and H. B. Shaffer. 2016. California amphibian and reptile species of special concern. University of California Press, Oakland, CA. 408 pp.

United States Fish & Wildlife Service. 2015. Endangered and threatened wildlife and plants; 90-day findings on 10 petitions. Federal Register 80: 19259–19263.

Vredenburg, V. T. 2004. Reversing introduced species effects: experimental removal of introduced fish leads to rapid recovery of a declining frog. Proceedings of the National Academy of Sciences 101: 7646–7650.

Walston, L. J. and S. J. Mullin. 2007. Responses of a pond-breeding amphibian community to the experimental removal of a predatory fish. The American Midland Naturalist 157: 63–73.

Winkler, J. D. and J. Van Buskirk. 2012. Influence of experimental venue on phenotype: multiple traits reveal multiple answers. Functional Ecology 26: 513–521.

